# Very Long-Chain Unsaturated Sphingolipids Mediate Oleate-Induced Rat β-Cell Proliferation

**DOI:** 10.1101/2021.07.16.452686

**Authors:** Anne-Laure Castell, Alexis Vivoli, Trevor S. Tippetts, Isabelle Robillard Frayne, Valentine S. Moullé, Matthieu Ruiz, Julien Ghislain, Christine Des Rosiers, William L. Holland, Scott A. Summers, Vincent Poitout

**Affiliations:** Montreal Diabetes Research Center, CRCHUM, Montréal, QC, Canada; Department of Medicine, Université de Montréal, Montréal, QC, Canada; Department of Nutrition and Integrative Physiology, University of Utah, Salt Lake City, UT, USA; Metabolomic Platform, Montreal Heart Institute Research Center, Montréal, QC, Canada; Department of Nutrition, Université de Montréal, Montréal, QC, Canada

**Author notes:** These authors contributed equally (Anne-Laure Castell, Alexis Vivoli). Corresponding author: Vincent Poitout, DVM, PhD, CRCHUM, 900 rue St Denis, Montréal, QC, H2X 0A9 - CANADA.

## Abstract

Fatty-acid (FA) signaling contributes to β-cell mass expansion in the face of nutrient excess, but the underlying mechanisms are poorly understood. Here we tested the hypothesis that sphingolipids, generated by the intracellular metabolism of FA, are implicated in the β-cell proliferative response to FA. Isolated rat islets were exposed to individual FA in the presence of 16.7 mM glucose for 48 h and the contribution of the *de novo* sphingolipid synthesis pathway was tested using the serine palmitoyltransferase inhibitor myriocin, the sphingosine kinase (SphK) inhibitor SKI II, or adenovirus-mediated knockdown of SphK, fatty-acid-elongase-1 (ELOVL1) and acyl-CoA-binding protein (ACBP). Wistar rat were infused with glucose and the lipid emulsion ClinOleic and received SKI II by gavage. B-cell proliferation was assessed by immunochemistry or flow cytometry. Sphingolipidomic analyses were performed by LC-MS/MS. Amongst the various FA tested, only oleate increased β-cell proliferation. Myriocin, SKI II, and SphK knockdown all decreased oleate-induced β-cell proliferation. Oleate exposure did not increase the total amount of sphingolipids but led to a specific rise in 24:1 species. Knockdown of ACBP or ELOVL1 inhibited oleate-induced β-cell proliferation. We conclude that unsaturated very long-chain sphingolipids produced from the available pool of C24:1 acyl-CoA mediate oleate-induced β-cell proliferation in rats.

---

The pancreatic β cell compensates for an increase in insulin demand, such as in obesity-associated insulin resistance, by expanding its functional mass (1). In rodents and possibly humans, the increase in β-cell mass is due at least in part to replication of existing β cells. Nutrients, including fatty acids (FA), play an important role in the β-cell proliferative response in both rodent and human islets (2,3). However, the underlying mechanisms of FA-induced β-cell proliferation remain elusive.

Previously we showed that 72-hour infusions of glucose and lipids in rats stimulate a compensatory increase in functional β-cell mass characterized by β-cell proliferation and insulin hypersecretion despite progressive β-cell dysfunction (4,5). These findings were reproduced *ex vivo* by showing that a mixture of 65% monounsaturated (MUFA), 20% polyunsaturated (PUFA), and 15% saturated (SFA) FA promotes β-cell proliferation in the presence of elevated glucose concentrations in isolated rat and human islets (3). Yet, the differential effects of individual FA species, and the intracellular FA metabolites mediating these effects, remain to be identified.

In the presence of elevated glucose concentrations, intracellular FA oxidation is inhibited and FA metabolism is preferentially shifted towards complex lipid synthesis, including acylglycerols and sphingolipids (6). This generates various intracellular signalling molecules that impact β-cell function and survival (7). Amongst these, sphingolipids including ceramides have emerged as important players in metabolic diseases (8,9). Ceramide accumulation is generally considered as detrimental, and mediates glucolipotoxicity-induced apoptosis and β-cell dysfunction (10–13). However, recent findings suggest that this view may be over simplistic as ceramides of different acyl chain lengths differentially affect glucose metabolism: whereas an increase in C16-C22 (long-chain; LC) ceramides is correlated with insulin resistance, a rise in C>22 (very long-chain; VLC) ceramides improves insulin signal transduction and protects mice from high-fat diet-induced obesity and glucose intolerance (14–16). Likewise, LC and VLC sphingolipids differentially affect cancer cell division and apoptosis (17–19). However, the differential effects of individual ceramide species and their metabolites on islet function has not been investigated.

Ceramides are produced through the *de novo* synthesis pathway in the endoplasmic reticulum following acylation of sphinganine by ceramide synthase (CerS) (Fig. 1A). As such, ceramides combine a sphingoid base and an amide-linked FA residue with a carbon chain length of C16-C24. Ceramides can be modified to produce sphingomyelins or glucosylceramides in the Golgi apparatus or metabolized to sphingosine which in turn is phosphorylated by sphingosine kinases (SphK) to generate sphingosine-1-phosphate (S1P). S1P promotes insulin secretion (20) and inhibits apoptosis in β-cell lines (21,22), and S1P administration promotes β-cell proliferation and inhibits apoptosis in diabetic mouse models (23). Furthermore, loss of SphK or S1P recycling enzymes leads to reduced β-cell mass and proliferation (24) and accelerates diabetes onset (25), respectively. Although these studies suggest a role of SphK/S1P in promoting β-cell function and mass, whether they contribute to β-cell proliferation in response to FA is currently unknown.

**Figure 1.**
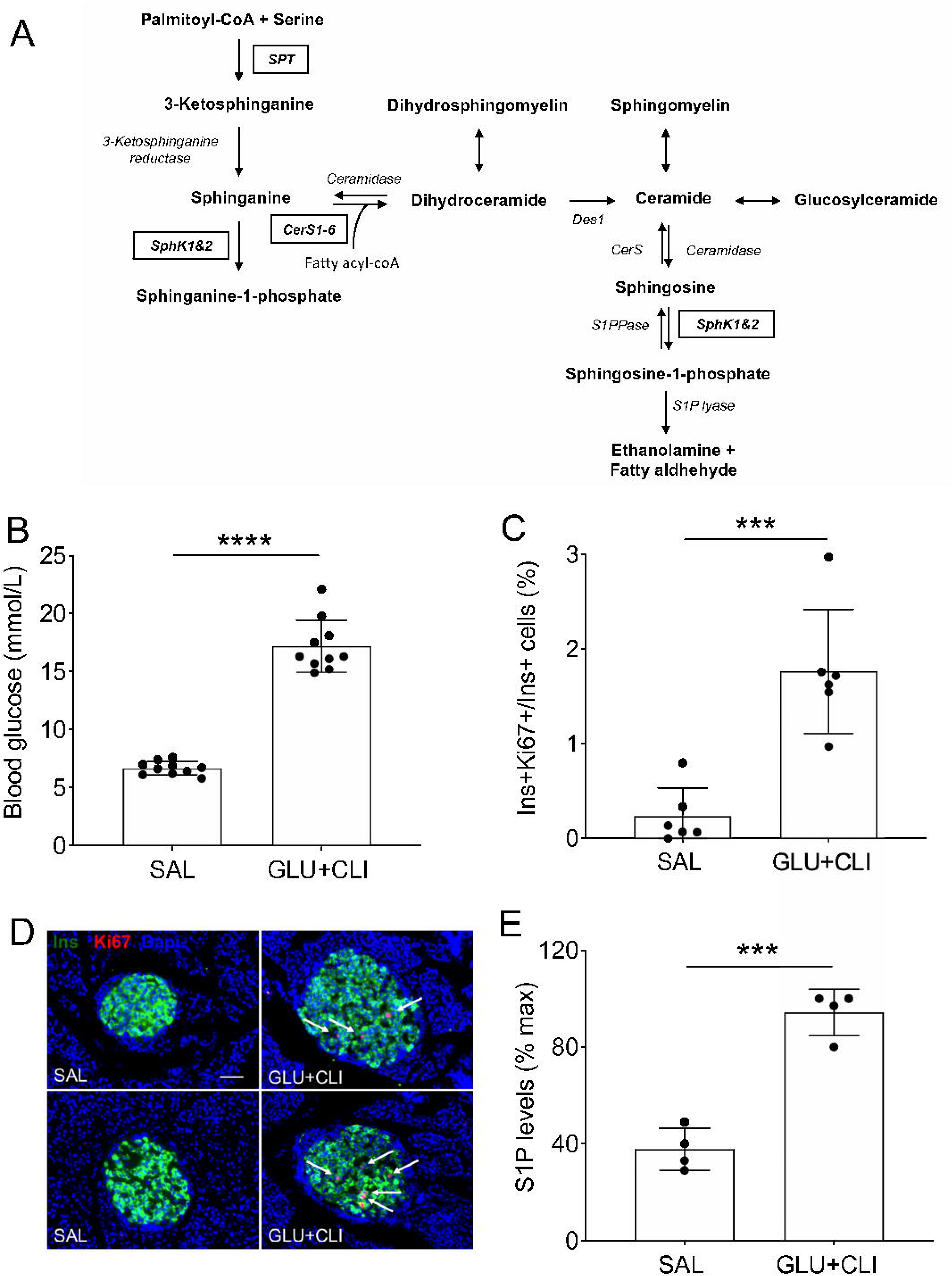
Infusions of GLU+CLI in Wistar rats increase β-cell proliferation and S1P levels in islets. (A) Biochemical pathway of sphingolipid metabolism. (B-E) 2-month-old Wistar rats were infused with saline (SAL) or glucose and ClinOleic (GLU+CLI) for 72 h (*n*=10). (B) Average blood glucose during the infusion. (C) β-cell proliferation was assessed by immunostaining for Ki67 and insulin (Ins) and presented as the percentage of Ki67^+^/Ins^+^ cells over total Ins^+^ cells (*n*=6). (D) Representative immunostaining for Ins (green), Ki67 (red) and nuclei (Dapi, blue) in pancreatic sections. Arrows show positive nuclei for Ki67. Scale bars, 50 μm. (E) S1P levels in islet extracts at the end of the infusion presented as percent of the maximum level. Data represent individual values and are expressed as mean +/− SEM. ***p<0.005, ****p<0.001 as compared with the SAL group by unpaired Student’s t-test. SPT, serine palmitoyl transferase; CerS, ceramide synthase; Des1, dihydroceramide desaturase 1; S1PPase, sphingosine-1-phosphate phosphatase; SphK, sphingosine kinase; S1P lyase, sphingosine-1-phosphate lyase.

The present study was designed to: 1- Identify the specific FA species that stimulate β-cell proliferation; 2- Assess the potential contribution of SphK/S1P to FA-induced β-cell proliferation; and 3-Determine the impact of sphingolipid acyl chain length and saturation on FA-induced β-cell proliferation.

## RESEARCH DESIGN AND METHODS

### Animals

All procedures were approved by the Institutional Committee for the Protection of Animals at the Centre de Recherche du Centre Hospitalier de l’Université de Montréal (CRCHUM). Two-month-old male Wistar rats (Charles River Laboratories, Saint-Constant, QC, Canada) were housed under controlled temperature on a 12-h light–dark cycle with free access to water and standard laboratory chow.

### Infusions

Rats underwent catheterisation of the jugular vein for systemic infusion of either saline (SAL) (0.9% w/v NaCl); or 70% (w/v) glucose plus 20% (w/v) ClinOleic (GLU+CLI) (an olive/soybean triacylglycerol emulsion composed of 65% MUFA/20% PUFA/15% SFA) as described (3,26). For *in vivo* studies, SKI II was dissolved in PEG400 (95%)/DMSO (5%) and administered by oral gavage once a day to GLU+CLI-infused rats at a final dose of 50 mg.kg^−1^. Control GLU+CLI-infused rats received SAL by oral gavage once a day. Measurement of plasma SKI II levels and immunostaining of pancreatic sections are described in Supplementary Materials.

### Islet Isolation, Culture and Adenoviral Infection

Islets were isolated by collagenase digestion and dextran density gradient centrifugation as described (27). Batches of 200 islets were cultured in RPMI-1640 with 10% (v/v) FBS or in MEM with 10% (v/v) dialysed FBS for 48 h in the presence of glucose (2.8 or 16.7 mM), a FA mixture mimicking the composition of CLI (MIX, 65% oleate, 20% linoleate and 15% palmitate) or FA alone as indicated in the figure legends. FA were complexed with BSA at a 5:1, 2.5:1 or 1.25:1 ratio (0.5, 0.25, 0.125 mM FA respectively) as indicated. To assess β-cell proliferation, 5-ethynyl-2’-deoxyuridine (EdU, 10 µM) was added throughout the culture period. Media were replaced daily. At the end of the treatment islets were processes for β-cell proliferation and apoptosis or sphingolipidomic analyses.

For adenoviral transduction, isolated islets were infected at an MOI of 1, 10 or 100 PFU/cell as described (28). 16 h later the medium was replaced with complete medium and cultured for an additional 24 h. Infected islets were cultured in the presence of glucose (2.8 or 16.7 mM) with or without oleate (0.5 mM) for 48 h with EdU (10 μM) prior to measuring β-cell proliferation of GFP^+^ and GFP^−^ cells. To assess knockdown efficiency, mRNA expression was measured by quantitative RT-PCR.

B-cell proliferation and apoptosis, quantitative RT-PCR and sphingolipidomic analyses are described in Supplementary Materials.

### Measurement of Sphingolipid Flux using Stable Isotopes

Batches of 200 islets were cultured in serine-deficient MEM with 10% (v/v) dialysed FBS for 2 h and then switched to MEM containing 0.4 mM L-serine-^13^C3,^15^N in the presence of 16.7 mM glucose and palmitate or oleate (0.5 mM) as indicated. After 16-h exposure, islets were washed with PBS and stored at −80° C. Sphingolipidomic analyses are described in Supplementary Materials.

### Statistical Analyses

Data are expressed as means ± SEM. Statistical analyses were performed using Student’s t test, one- or two-way ANOVA with Tukey’s or Dunnett’s post hoc test adjustment for multiple comparisons, as appropriate, using GraphPad InStat (GraphPad Prism 8 Software, San Diego, CA). p <0.05 was considered significant.

### Data and Resource Availability

All data generated or analyzed during this study are included in the published article (and its online supplementary files). No applicable resources were generated or analyzed during the current study.

## RESULTS

### Nutrient infusions in Wistar rats increase S1P concentrations in islets

Given the reported effects of S1P on β-cell proliferation (23), we first assessed the effect of nutrient excess on S1P concentrations in islets using our rat model of GLU+CLI infusion (3,4). Wistar rats were infused with SAL or GLU+CLI for 72 h. GLU+CLI infusion increased blood glucose in the target range (≈15 mmol/L) (Fig. 1B) and, consistent with previous studies (3), led to an increase in β-cell proliferation as assessed by the percentage of Ki67^+^/Ins^+^ cells (Fig. 1C and D). S1P concentrations were increased in islet extracts of GLU+CLI-infused rats compared to the SAL-infused group (Fig. 1E), suggesting that GLU+CLI infusion enhances the flux through the *de novo* ceramide synthesis pathway in islets *in vivo.*

### Oleate selectively stimulates β-cell proliferation in rat islets

As CLI is composed of MUFA, PUFA and SFA we measured the effect of each type of FA on β-cell proliferation in isolated rat islets. Islets were exposed to a FA mixture, mimicking the composition of CLI (MIX), or to palmitate (SFA, C16:0), oleate (MUFA, C18:1) or linoleate (PUFA, C18:2) in the presence of 16.7 mM glucose for 48 h. B-cell proliferation was assessed by immunostaining of islet sections or by flow cytometry. Immunohistochemistry revealed that the FA mixture increased β-cell proliferation as expected (Fig. 2A and B). The increase in β-cell proliferation in response to oleate was comparable to the MIX, but palmitate and linoleate had no effect. Likewise, oleate but not palmitate dose-dependently increased β-cell proliferation as assessed by flow cytometry (Fig. 2C-E). Neither oleate nor palmitate affected α-cell proliferation (Fig. 2F). A significant increase in β-cell apoptosis was detected in response to palmitate, but not oleate (Supplementary Fig. 1A). The other MUFA palmitoleate (C16:1) did not significantly increase β-cell proliferation (Supplementary Fig. 1B). Overall, these results show that amongst the various FA tested, only oleate stimulates β-cell proliferation, suggesting that the carbon chain length and the position and degree of saturation are important in this process.

**Figure 2.**
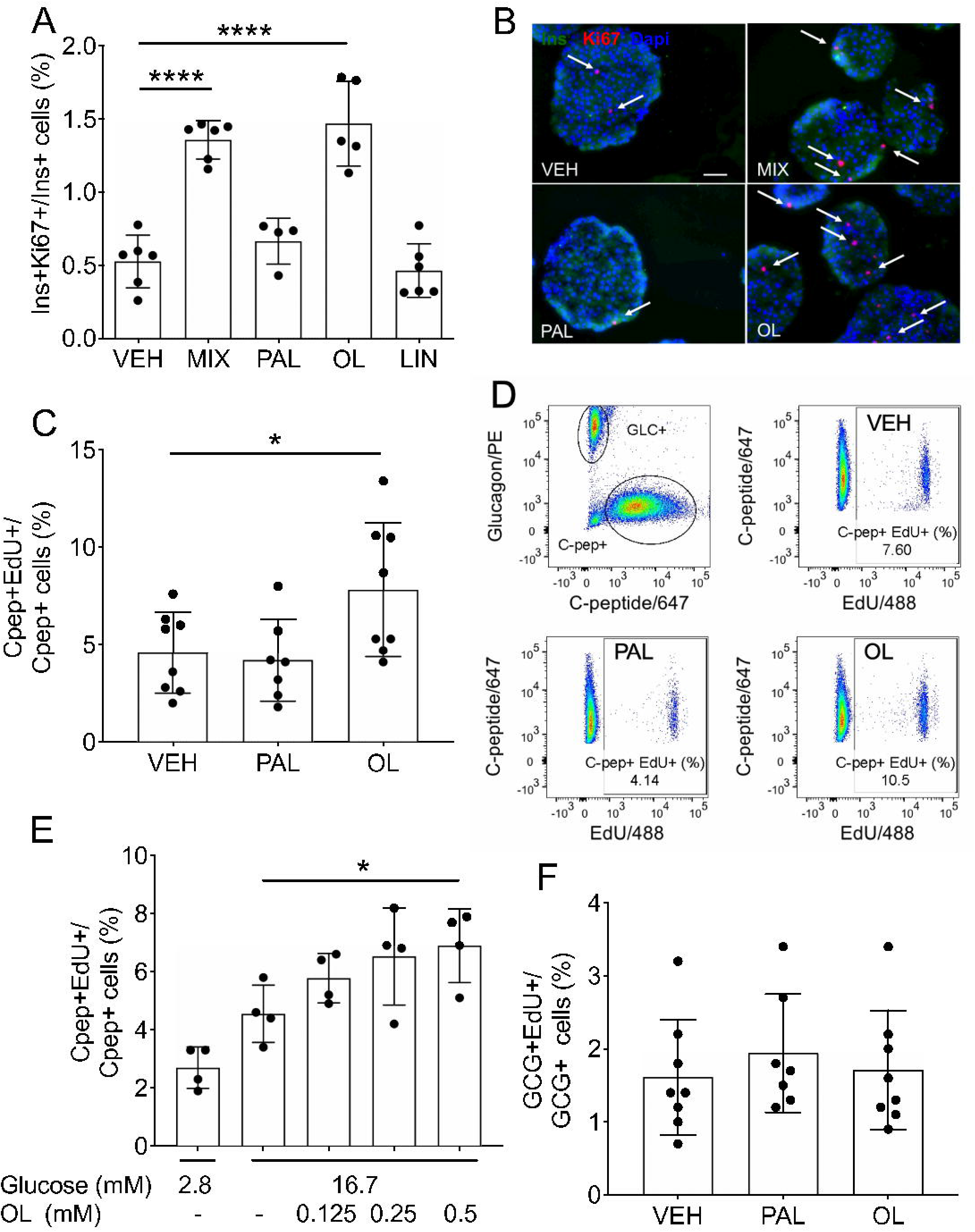
Oleate promotes β-cell proliferation in isolated rat islets. Isolated rat islets were exposed to 16.7 mM glucose and a mixture of fatty acid (MIX), palmitate (PAL), oleate (OL) or linoleate (LIN) at 0.5 mM, or vehicle (VEH) (A-D, F) or 2.8 or 16.7 mM glucose and OL (0.125-0.5 mM) (E) for 48 h, as indicated. B-cell proliferation was assessed by immunostaining for Ki67 and insulin (Ins) (A, B) and presented as the percentage of Ki67^+^/Ins^+^ cells over total Ins^+^ cells (A), and by flow cytometry following staining for EdU and C-peptide (Cpep) (C-E) and presented as the percentage of EdU^+^/Cpep^+^ cells over total Cpep^+^ cells (C, E). A-cell proliferation was assessed by flow cytometry following staining for EdU and glucagon (GCG) and presented as the percentage of EdU^+^/GCG^+^ cells over total GCG^+^ cells (F). (B) Representative images of immunostaining for Ins (green), Ki67 (red) and nuclei (Dapi, blue) in pancreatic sections. Arrows show cells positive for Ki67 and insulin. Scale bars, 50 μm. (D) Representative plots showing gates used to select Cpep^+^ and GCG^+^ cells and EdU^+^/Cpep^+^ cells in VEH, PAL and OL conditions. Data represent individual values and are expressed as mean + /− SEM (*n*=4-8). *p<0.05, ****p<0.001 by one-way ANOVA with Dunnett’s (A, C, F versus VEH condition) or Tukey’s (E) multiple comparisons tests.

### Oleate-induced β-cell proliferation requires *de novo* sphingolipid synthesis and is blocked upon inhibition of SphK

Our findings so far suggest that 1-β-cell proliferation is associated with *de novo* sphingolipid synthesis in vivo, and 2-FA stimulation of β-cell proliferation is restricted to oleate. These two observations are seemingly contradictory, since palmitate, but not oleate, is the preferential FA substrate for the first step of *de novo* sphingolipid synthesis. To confirm the implication of this pathway, we first blocked that step with the serine palmitoyl transferase (SPT) inhibitor Myriocin. Myriocin dose-dependently decreased oleate-induced β-cell proliferation (Fig. 3A and B). Second, we depleted the culture medium of L-serine, the other substrate of the SPT reaction. Islets cultured in L-serine-deficient medium were unresponsive to oleate, and the addition of L-serine restored the β-cell proliferative response (Fig. 3C).

**Figure 3.**
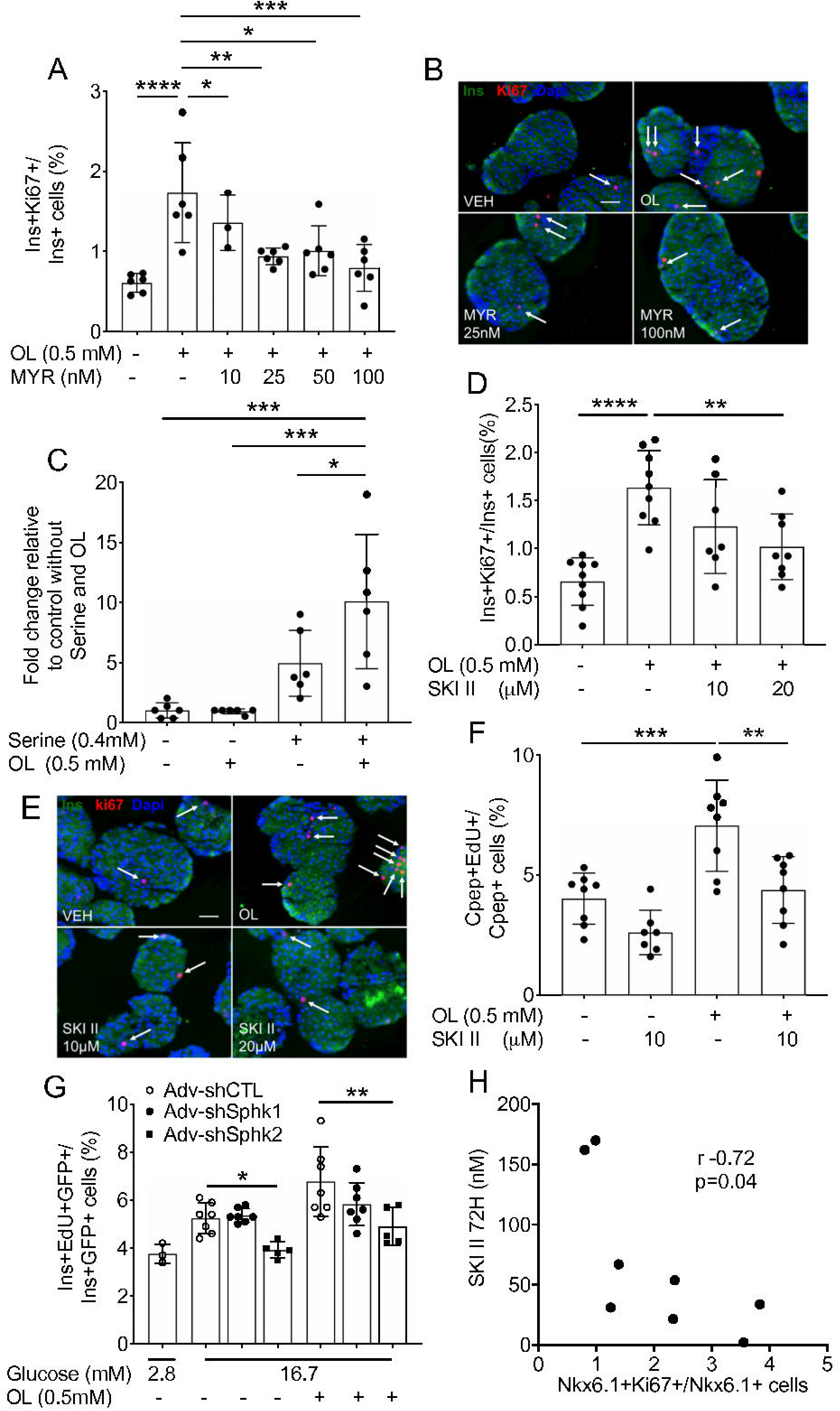
Blockade of *de novo* sphingolipid synthesis and inhibition of SphK decrease oleate-induced β-cell proliferation in isolated rat islets. (A, B, D-G) Isolated rat islets were exposed to oleate (OL; 0.5 mM) or vehicle (VEH) in the presence of 16.7 mM glucose and the serine palmitoyl transferase inhibitor myriocin (MYR; 10 to 100 nM) (*n*=3-6) (A, B) or the sphingosine kinase (SphK) inhibitor (SKI II; 10 or 20 μM) (*n*=7-9) (D-F), or following adenoviral-mediated knockdown of SphK1 (Adv-shSphk1) or SphK2 (Adv-shSphk1) or control adenovirus (Adv-shCTL) (*n*=3-7) (G) as indicated. (C) Isolated rat islets were cultured in L-serine deficient MEM and exposed to 16.7 mM glucose and OL (0.5 mM) with or without L-serine (0.4 mM) for 48 h. B-cell proliferation was assessed by immunostaining for Ki67 and insulin (Ins) (A, B, D, E) and presented as the percentage of Ki67^+^/Ins^+^ cells over total Ins^+^ cells (A, D); by flow cytometry following staining for EdU and C-peptide (Cpep) or insulin (Ins) (C-E) and presented as the percentage of EdU^+^/Cpep^+^ over total Cpep^+^ (C, F), as fold-change over the control condition without L-serine and OL (C), or as EdU^+^/Ins^+^/GFP^+^ over total Ins^+^/GFP^+^ cells (G). (B, E) Representative images of Ins (green), Ki67 (red), nuclei (Dapi, blue) staining. Arrows show positive nuclei for Ki67. Scale bars, 50 μm. (H) Correlation between β-cell proliferation expressed as the percentage of Nkx6.1^+^/Ki67^+^ cells over total Nkx6.1^+^ cells and plasma SKI II levels (nM) after 72 h infusion of GLU+CLI and SKI II (50mg/kg/d) administration (*n*=8). Data represent individual values and are expressed as mean +/− SEM. *p<0.05, **p<0.01, ***p<0.005, ****p<0.001 by one-way ANOVA with Tukey’s (A, C, D, F) multiple comparisons test and Mixed effect analysis with Dunett’s test for multiple comparisons (G, versus Adv-shCTL conditions) or Pearson correlation (H).

We then asked whether SphK/S1P contributes to oleate-induced β-cell proliferation. As rat islets express two SphK isoforms, SphK1 and SphK2 (29), we blocked total SphK activity with the inhibitor SKI II. SKI II inhibited oleate-stimulated β-cell proliferation as assessed by immunohistochemistry (Fig. 3D and E) and flow cytometry (Fig. 3F), but not the response to 16.7 mM glucose alone (Fig. 3F). Treatment with SKI II was not associated with detectable apoptosis (Supplementary Fig. 1C).

To substantiate the findings obtained with SKI II, we infected isolated rat islets with adenoviruses expressing shRNA against SphK1 and SphK2. To titrate the viral load and exclude non-specific effects of the infection on β-cell proliferation, we first infected islets for 48 h with increasing titers of the GFP-expressing control adenovirus and quantified GFP^+^ β cells by flow cytometry. Following infection with 1, 10 and 100 PFU/cell, approximately 10, 25 and 40% of Cpep^+^ cells were GFP^+^, respectively, and these percentages were similar across culture conditions (Supplementary Fig. 2A). Surprisingly, adenoviral infection at 10 and 100 PFU/cell inhibited oleate stimulation of β-cell proliferation without affecting the response to glucose (Supplementary Fig. 2B). Despite this limitation, we choose to use 100 PFU/cell for optimal gene knockdown. Compared to Adv-shCTL, infection with Adv-shSphK1 and Adv-shSphK2 led to a significant reduction in SphK1 and SphK2 RNA levels, respectively, in GFP^+^ cells at 48 h post-infection (Supplementary Fig. 2C and D). Knockdown of SphK2, but not of SphK1, led to a significant decrease in the proliferative response to glucose and oleate (Fig. 3G), consistent with the reported weaker expression of SphK1 compared to SphK2 in β cells (30).

To investigate the role of SphK in nutrient-induced β-cell proliferation *in vivo* we administered SKI II (50mg/kg/d) by oral gavage to 72-h GLU+CLI-infused rats. SKI II treatment did not affect target blood glucose levels (Supplementary Fig. 3A) or glucose infusion rates (GIR, mg/kg/min) in GLU+CLI-infused animals (Supplementary Fig. 3B). Whole body and pancreas weights at the end of the infusion were reduced in GLU+CLI-infused animals compared to the SAL-infused group but not affected by SKI II treatment (Supplementary Fig. 3C and D). Plasma SKI II levels increased overtime but were highly variable between animals by the end of the infusion (Supplementary Fig. 3E). Accordingly, the effect of SKI II on GLU+CLI-induced β-cell proliferation, as assessed by immunostaining for Nkx6.1 and Ki67, was also highly variable and therefore not significantly different from the control group (Supplementary Fig. 3F). Hence, we plotted plasma SKI II levels at 72 h against β-cell proliferation in individual animals (Fig. 3H) and noted a significant inverse correlation (r=−0.72), indicating that increased plasma SKI II levels are associated with a decrease in GLU+CLI-induced β-cell proliferation.

Taken together, these data suggest that *de novo* sphingolipid synthesis and SphK activity are necessary for the β-cell proliferative response to oleate.

### S1P and Sa1P do not contribute to oleate-induced β-cell proliferation

We then asked whether the products of SphK, S1P and sphinganine-1-phosphate (Sa1P), mediate the effect of oleate on β-cell proliferation. Surprisingly, S1P levels increased in islets following 48 h exposure to palmitate, but not oleate (Fig. 4A). SKI II, at a concentration shown to inhibit β-cell proliferation (10 μM; Fig. 3), did not significantly diminish S1P or Sa1P levels in the presence of oleate or 16.7 mM glucose alone (Fig. 4B and C). Finally, blockade of S1P lyase, the enzyme that catalyzes the irreversible degradation of S1P, with THI did not affect β-cell proliferation in response to oleate or 16.7 mM glucose alone (Fig. 4D). These results suggest that the sphingolipid metabolites S1P and Sa1P are not involved in oleate-induced β-cell proliferation.

**Figure 4.**
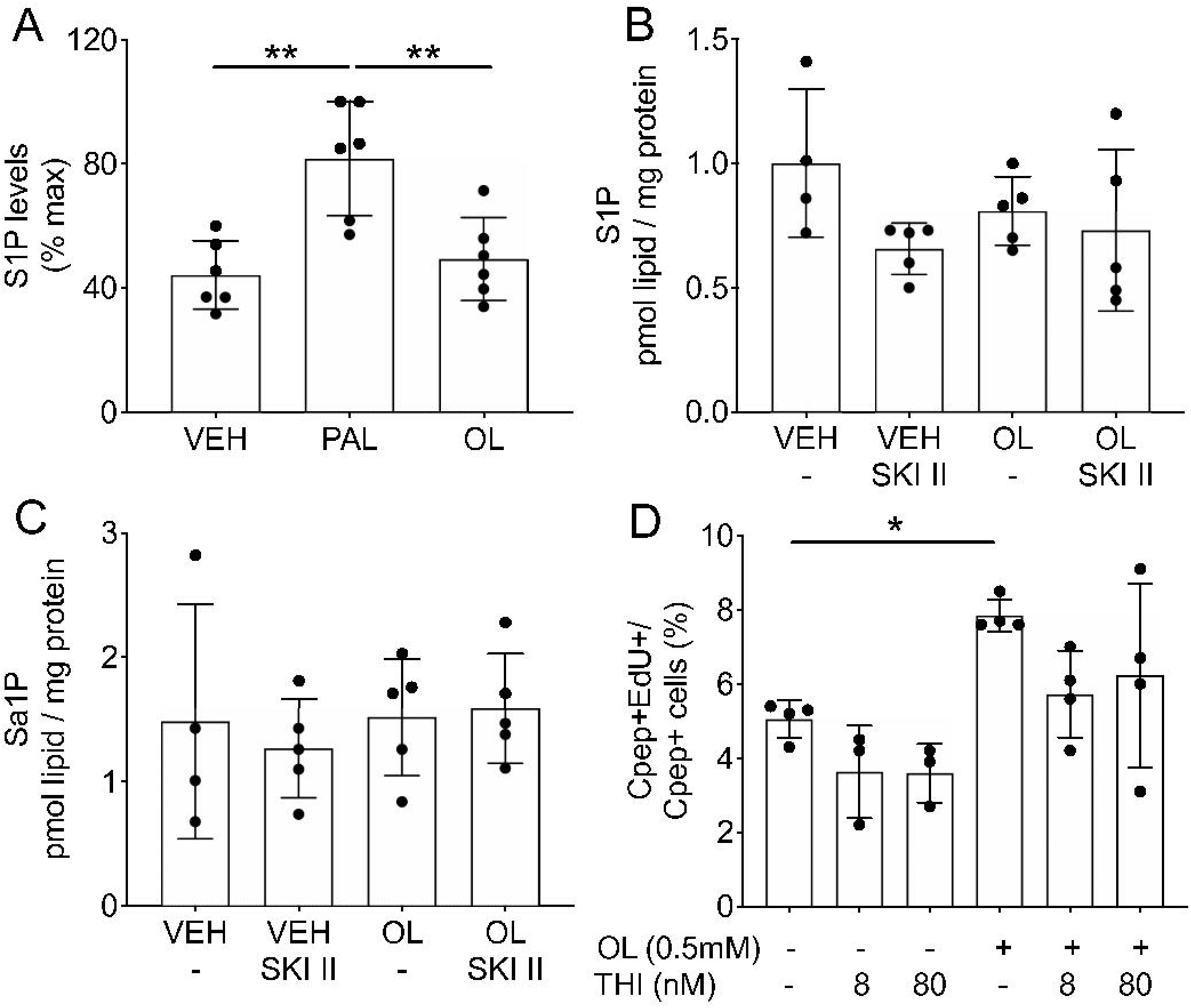
Sphingolipid metabolites S1P or Sa1P do not contribute to oleate-induced β-cell proliferation in isolated rat islets. (A) Relative sphingosine-1-phosphate (S1P) levels in islet extracts after 48 h exposure to palmitate (PAL, 0.5 mM), oleate (OL, 0.5 mM) or vehicle (VEH) (*n*=6). (B, C) S1P and dihydroS1P (Sa1P) levels in islets extracts after 48 h exposure to OL (0.5 mM) in the presence of 16.7 mM glucose with or without SKI II (10 μM) (*n*=5). (D) Islets were exposed to OL (0.5 mM) in the presence of 16.7 mM glucose and the S1P lyase inhibitor THI (8 or 80 nM) for 48 h (*n*=3-4) and β-cell proliferation assessed by flow cytometry following staining for EdU and C-peptide (Cpep) and presented as the percentage of EdU^+^/Cpep^+^ over total Cpep^+^. Data represent individual values and are expressed as mean +/− SEM. *p<0.05, ***p<0.005 using one-way ANOVA with Dunnett’s (D, versus VEH condition) or Tukey’s (A-C) multiple comparisons tests.

### Palmitate and oleate differentially affect the sphingolipid profile in islets

Although palmitate is the preferential FA for the sphingoid base of ceramides and complex sphingolipids, both oleate and palmitate are substrates for the amino-linked FA portion (Fig. 1A). FA elongases (ELOVL) extend FA chain length to C16-C24 depending on the substrate. Hence, oleate and palmitate might affect islet sphingolipid synthesis at the level of the amino-linked FA chain length rather than total levels of sphingolipid classes. To test for this possibility, we performed a comprehensive and untargeted analysis of sphingolipids by LC-MS/MS on lipid extracts following islet treatment with palmitate or oleate for 48 h (Fig. 5A). Principal component analysis (PCA) revealed major differences in steady-state lipid profiles between vehicle-, palmitate-, and oleate-treated islets (Fig. 5B). Whereas palmitate increased the level of saturated LC and VLC ceramides (C16:0-C24:0), oleate only increased the unsaturated VLC ceramides (C24:1) (Fig. 5C). In addition, palmitate increased C22:0 and C24:0 but decreased C24:1 sphingomyelins, while oleate diminished C16:0-C24:0 and increased C24:1 sphingomyelins (Fig. 5D). Similarly, palmitate increased C16:0 and C22:0 but decreased C24:1 glucosylceramides, whereas oleate diminished C22:0 and increased C24:1 glucosylceramides (Fig. 5E). In a complementary analysis performed under similar conditions we observed that the total amount of islet sphingolipids was not affected by either palmitate or oleate treatment (Supplementary Fig. 4A). However, as with the previous analyses, amino-linked FA chain length was altered: palmitate increased several saturated LC and VLC sphingolipid species while oleate decreased these saturated sphingolipids and increased unsaturated VLC (C24:1) sphingolipid species (Supplementary Fig. 4B-F).

**Figure 5.**
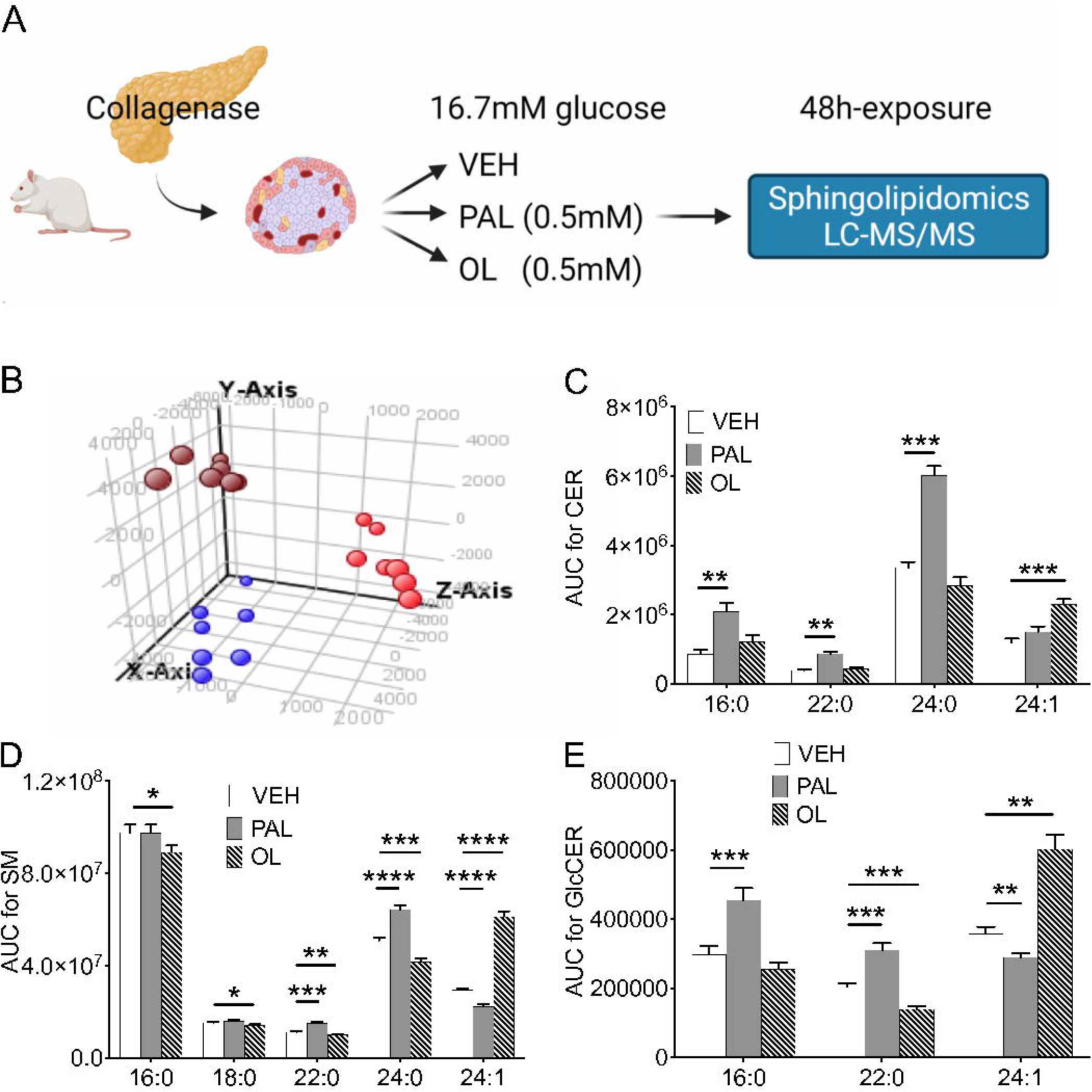
Palmitate and oleate differentially affect sphingolipid FA side-chain length and saturation. (A) Lipidomic analyses were performed by LC-MS/MS on lipid extracts following 48 h-exposure of islets to palmitate (PAL, 0.5mM), oleate (OL, 0.5mM) or vehicle (VEH) in the presence of 16.7 mM glucose (*n*=7). Image created with Biorender.com. (B) 3D PCA score plot of lipids comparing the vehicle (VEH) (red), PAL (brown) and OL (blue) treatment conditions. (C-E) Area under the curve (AUC, normalized to cyclic loess) of ceramides (CER) (C), sphingomyelins (SM) (D) and glucosylceramides (GlcCER) (E) grouped according to amino-linked FA chain length and saturation. Data are expressed as mean +/− SEM. *p<0.05, **p<0.01, ***p<0.005, ****p<0.001 using two-way ANOVA with Dunnett’s multiple comparisons test as compared with the VEH condition.

To confirm that palmitate and oleate stimulate *de novo* sphingolipid synthesis we performed stable-isotope labelling of sphingolipids. Islets were cultured in the presence of L-serine-^13^C_3_, ^15^N, and sphingolipids were analyzed following 16 h-exposure to palmitate or oleate (Fig. 6A). Palmitate, but not oleate, increased total isotope-labelled dihydroceramides and ceramides (Fig. 6B and C). Importantly, palmitate increased C16:0-C24:0, whereas oleate increased C24:1 isotope-labelled dihydroceramides and ceramides (Fig. 6D and E).

**Figure 6.**
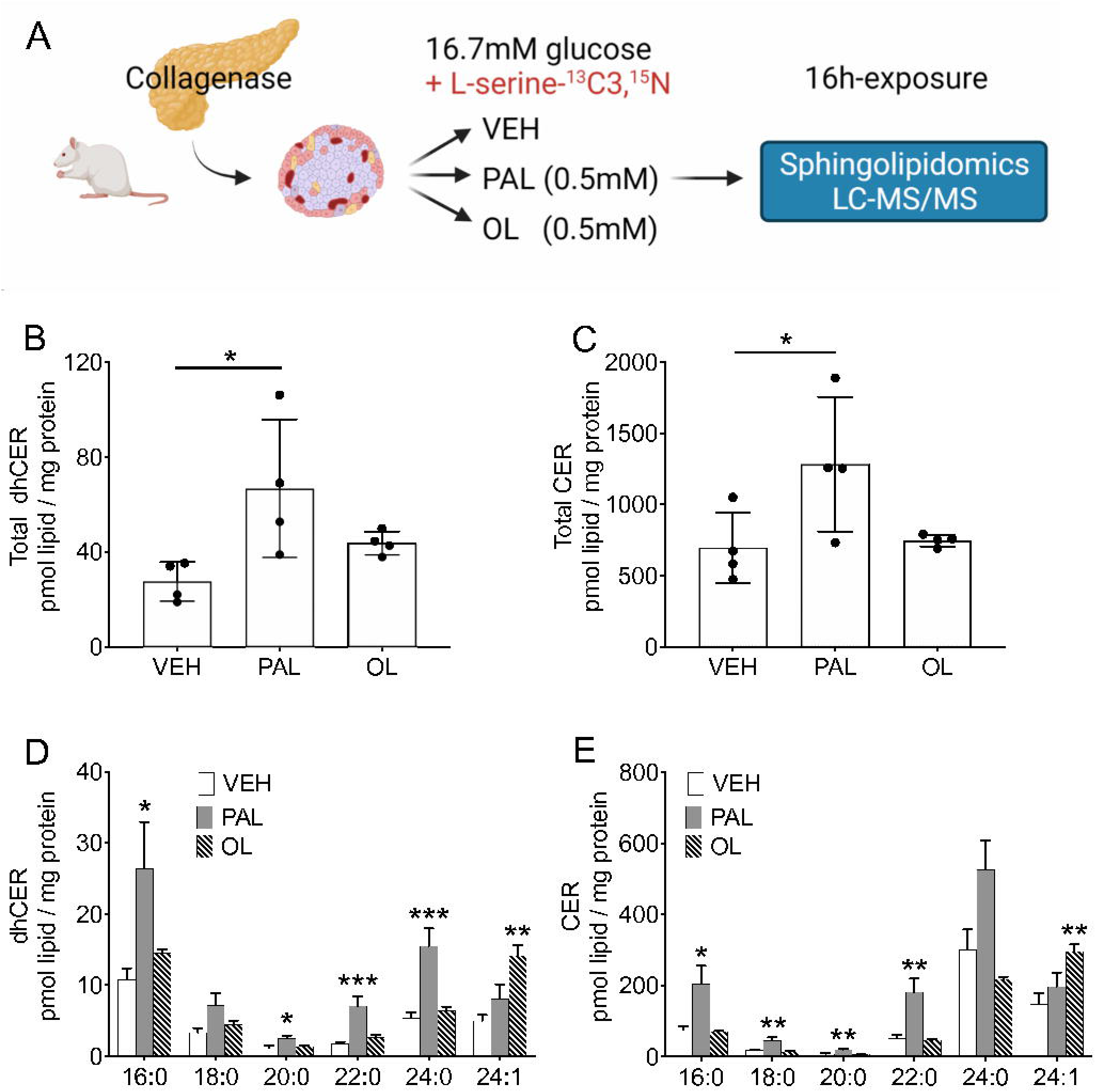
Palmitate and oleate regulate sphingolipid FA side-chain characteristics through the *de novo* pathway. (A) Lipidomic analyses were performed by LC-MS/MS on lipid extracts following 16 h-exposure of isolated islets to palmitate (PAL, 0.5 mM), oleate (OL, 0.5 mM) or vehicle (VEH) in the presence of 16.7 mM glucose and stable-isotope labelled serine (0.4 mM) (*n*=4). Image created with Biorender.com. (B-E) Normalized levels (pmol lipid/mg protein) of total dihydroceramides (dhCER) (B) and ceramides (CER) (C) and specific dhCER (D) and CER (E) grouped according to amino-linked FA chain length and saturation. Data are expressed as mean +/− SEM. *p<0.05, **p<0.01, ***p<0.005 using one-way ANOVA with Dunnetts’s multiple comparisons test as compared to the VEH condition.

Taken together, these data indicate that palmitate and oleate impact amino-linked FA characteristics through *de novo* sphingolipid synthesis and that oleate exposure leads to increased synthesis of C24:1 species without an increase in total ceramide levels.

### SKI II alters levels of sphingolipid species in oleate-treated islets

As inhibition of SphK with SKI II blocked oleate stimulation of β-cell proliferation without significantly affecting S1P or Sa1P levels, we asked whether this effect could be mediated by changes in other sphingolipid species. We analyzed lipid extracts following islet exposure to oleate in the presence of SKI II. Addition of SKI II significantly increased the total amount of ceramides in oleate-treated islets (Fig. 7A). Although not significant, there was a trend towards an increase in total dihydroceramides, dihydrosphingomyelins and sphingomyelins in the presence of SKI II. Interestingly, these changes were often associated with an increase in sphingolipids with specific acyl chains. Hence, C24:1 dihydroceramides (Fig. 7B) and 16:0 ceramide and dihydrosphingomyelins (Fig. 7C and D), but not sphingomyelin or glucosylceramide species (Fig. 7E and F), were augmented in response to SKI II exposure. Overall, these data indicate that the inhibition of β-cell proliferation by SKI II is associated with an increase in several C16:0 sphingolipid species.

**Figure 7.**
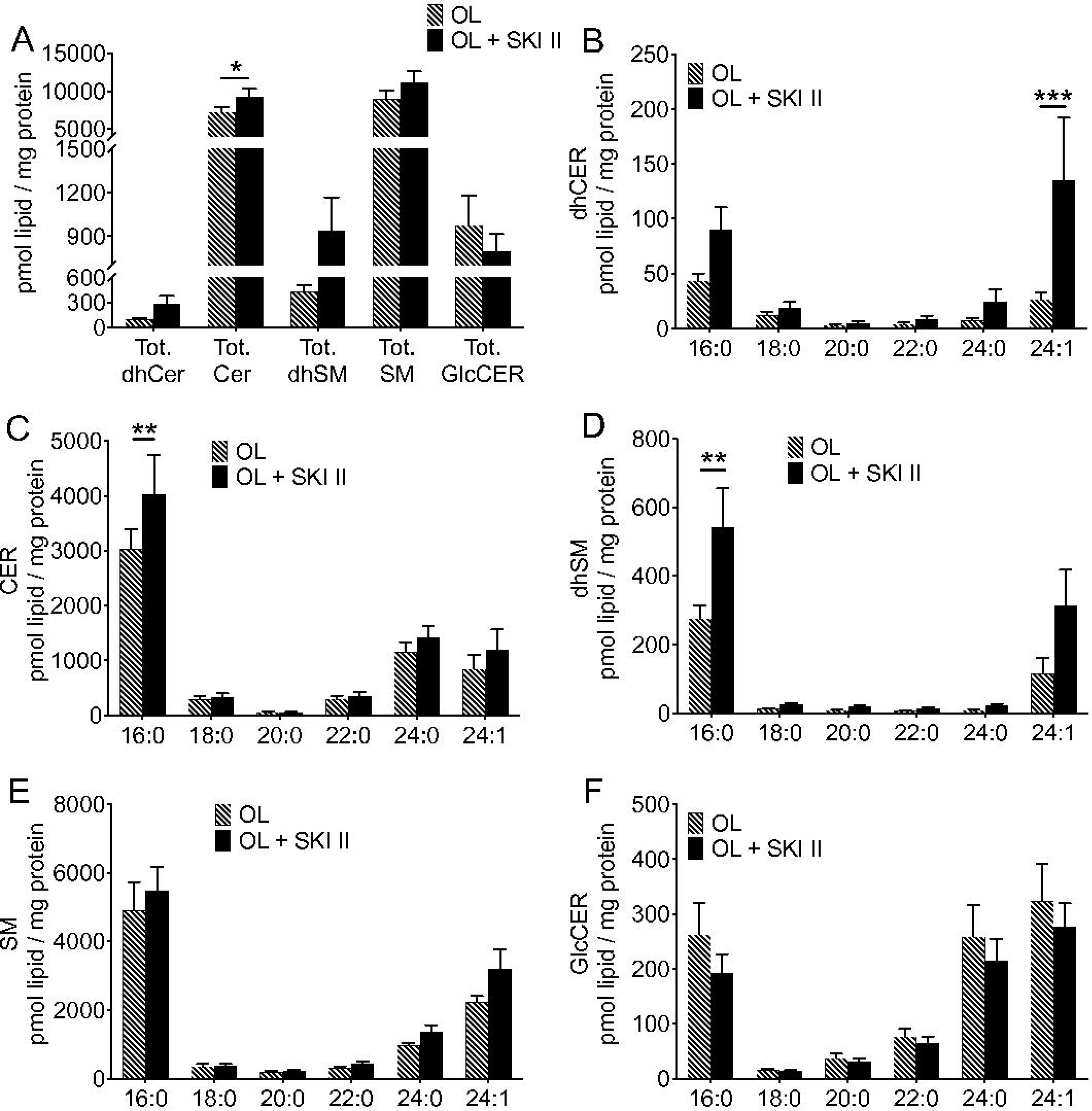
SKI II increases the levels of several sphingolipid species in oleate-treated islets. Lipidomic analyses were performed by LC-MS/MS on lipid extracts following 48 h-exposure of isolated islets to oleate (OL, 0.5 mM) in the presence of 16.7 mM glucose with or without the sphingosine kinase inhibitor, SKI II (10 μM). (A-F) Normalized levels (pmol lipid/mg protein) of total sphingolipids (A) or dihydroceramides (dhCER) (B), ceramides (CER) (C), dihydrosphingomyelins (dhSM) (D), sphingomyelins (SM) (E) and glucosylceramides (GlcCER) (F) grouped according to amino-linked FA chain length and saturation. Data are expressed as mean +/− SEM (*n*=5-8). *p<0.05, **p<0.01, ***p<0.005 using two-way ANOVA with Tukey’s multiple comparisons test.

### Blockade of VLC sphingolipid synthesis inhibits oleate-induced β-cell proliferation

Our results so far suggest that β-cell proliferation is regulated by the relative balance between individual sphingolipid species: conditions associated with an increase in C24:1 species, such as in the presence of oleate, are stimulatory, whereas those associated with an increase in C16:0 sphingolipids, such as in the presence of palmitate or SKI II, are neutral or inhibitory. To further ascertain the involvement of C24:1 sphingolipids in β-cell proliferation induced by oleate, we knocked down ELOVL1 or acyl-CoA binding protein (ACBP) in islets exposed to oleate. Knockdown of ACBP is expected to reduce the available pool of LC acyl-CoA for elongation by ELOVL1 and consequently decrease the synthesis of VLC sphingolipids (31–33). Infection of islets with Adv-shELOVL1 or Adv-shACBP significantly diminished their respective mRNA levels (Supplementary Fig. 5A and B). Knockdown of ELOVL1 completely blocked the ability of glucose and oleate to stimulate β-cell proliferation, and knockdown of ACBP significantly decreased the response to oleate, in GFP^+^ (Fig. 8A and B), but not GFP^−^ (Supplementary Fig. 5C and D) cells. These data indicate that the synthesis of VLC sphingolipids via ELOVL1 is necessary for oleate-induced β-cell proliferation.

**Figure 8.**
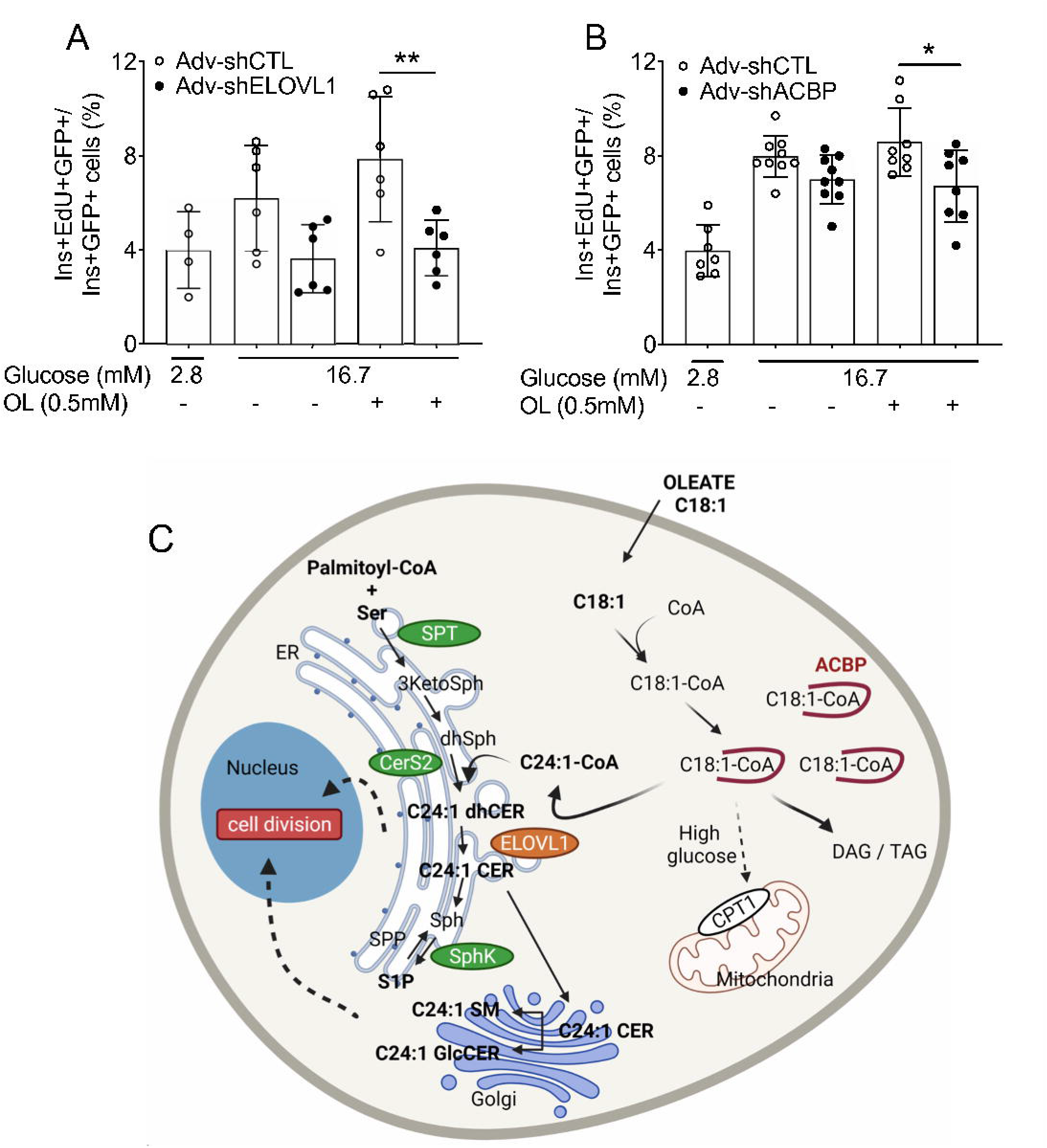
Knockdown of ELOVL1 and ACBP inhibits oleate induced β-cell proliferation in isolated rat islets. Isolated rat islets were infected with Adv-shCTL, Adv-shELOVL1 (A) or Adv-shACBP (B) and exposed to 2.8 or 16.7 mM glucose with or without oleate (OL, 0.5 mM) as indicated. Proliferation of infected (GFP^+^) β cells was assessed by flow cytometry following staining for EdU and insulin (Ins) and presented as the percentage of EdU^+^/Ins^+^/GFP^+^ over total Ins^+^/GFP^+^ cells. Data are presented as fold-change over the control condition (Adv-shCTL). Data represent individual values and are expressed as mean +/− SEM. *p<0.05, **p<0.01 using mixed-effect analysis with Sidak’s multiple comparisons test as compared with Adv-shCTL condition (A, B). (C) Proposed mechanism whereby the oleate/very long chain sphingolipid axis controls β-cell proliferation. In high glucose condition when FA oxidation is inhibited, oleate transformed in C18:1-CoA in the cell is stabilized by ACBP thus favouring its elongation to C24:1 by ELOVL1 and the synthesis of C24:1 sphingolipids after acylation by CerS2. The production of C24:1 sphingolipids leads to an increase in β-cell proliferation. Image created with Biorender.com.

## DISCUSSION

The objectives of this study were to identify the specific FA species stimulating rat β-cell proliferation and to investigate the underlying mechanisms. We showed that among several FA tested, only oleate potentiates β-cell proliferation in the presence of elevated glucose concentrations. This was unexpectedly associated with increased *de novo* sphingolipid synthesis which did not lead to higher S1P and Sa1P levels but to a selective increase in unsaturated VLC (C24:1) sphingolipids. Importantly, blocking VLC (C24:1) sphingolipid synthesis inhibited oleate-induced β-cell proliferation. Overall, these results identify a key role for a FA-CoA-C24:1 sphingolipid axis in the β-cell proliferative response to nutrients.

Infusion of GLU+CLI in rats increased β-cell proliferation as did a lipid mixture mimicking the FA composition of CLI in isolated rat islets. Only the MUFA oleate (C18:1) reproduced the effect of the lipid mixture on β-cell proliferation. These data are consistent with previous studies demonstrating a pro-proliferative role of a MUFA-enriched emulsion *in vivo* and oleate *ex vivo* in rat β cells (3,34). Although previous studies in mice reported an inhibition of glucose-induced β-cell proliferation by palmitate and linoleate (35), neither FA affected β-cell proliferation in rat islets *ex vivo* in the present study. However, palmitate did increase β-cell apoptosis, as shown previously (36). Collectively these findings highlight the key role of chain length and degree of saturation in FA-induced β-cell proliferation, as reported previously for FA-induced β-cell dysfunction (13,37).

Inhibition of β-cell proliferation in the presence of the SPT inhibitor or following depletion of L-serine demonstrated that *de novo* sphingolipid synthesis is implicated in the response to oleate. Our observations that pharmacological inhibition or knockdown of SphK inhibited oleate-induced β-cell proliferation were suggestive of a role for the SphK metabolites S1P and Sa1P, as shown in cancer cell lines (38). Surprisingly however, oleate did not increase S1P or Sa1P levels in isolated islets, suggesting that the rise in islet S1P levels following GLU+CLI infusion was likely due to the presence of palmitate in the ClinOleic emulsion. In addition, although SKI II decreased oleate-induced β-cell proliferation, it did not alter S1P or Sa1P levels in isolated islets. Furthermore, blocking S1P degradation with the S1P lyase inhibitor had no effect on β-cell proliferation. This led us to conclude that oleate-induced β-cell proliferation is not mediated by S1P or Sa1P. Although SphK/S1P exerts a critical role in β-cell survival in response to lipotoxic or cytokine stress *in vitro* (21,39,40) and in response to high-fat feeding *in vivo* (25), the role of S1P in β-cell proliferation is unresolved. Whereas β-cell proliferation increases in streptozotocin-induced diabetic mice following S1P administration (23), inhibiting S1P metabolism in S1P phosphatase 2 knockout mice reduces β-cell proliferation in response to high-fat feeding (24). Furthermore, exogenous S1P compromises insulin-induced β-cell proliferation in MIN6 cells (41).

In cancer cell lines SKI II inhibits not only SphK activity but also dihydroceramide desaturase 1 activity leading to a loss of S1P and an accumulation of dihydroceramide and dihydrosphingomyelin (42,43). Although no significant changes in S1P were detected in oleate-treated rat islets exposed to SKI II, an increase in total ceramides, specifically C16:0 ceramides, as well as C24:1 dihydroceramides and C16:0 dihydrosphingomyelins was observed. Given that ceramide levels are at least an order of magnitude higher than dihydroceramides and dihydrosphingomyelins, changes in ceramide levels are likely to be more biologically significant. The decrease in oleate-induced β-cell proliferation following depletion of SphK activity might be due to a change in the relative balance of specific sphingolipid species, notably the C16:0/C24:1 ratio. In support of this possibility, whereas S1P promotes cell survival and proliferation, ceramides mediate apoptosis and growth inhibition. Antagonistic effects between sphingolipid species has led to the concept of the “sphingolipid rheostat”, highlighting a critical role of the dynamic balance between ceramides and S1P in the control of cell biology (44). Furthermore, N-acyl chain length significantly impacts the biological activities of ceramides (18,45). Whereas CerS2^−/−^ mice present glucose intolerance associated with an increase in C16 and a decrease in C24 ceramides in the liver (16,46), loss of CerS6 protects from diet-induced obesity and insulin resistance by decreasing C16 ceramides (15,47). In the β cell, C14:0-C24:0 ceramides are associated with palmitate-mediated apoptosis and inhibition of insulin secretion (10,11). Similarly, in cancer cells upregulation of CerS4 and 6 leads to an increase in C16:0-C20:0 ceramides and inhibition of cell proliferation (17). Further studies will be required to determine whether saturated LC and VLC ceramide species negatively regulate β-cell proliferation.

In the presence of high glucose concentrations, oleoyl-CoA is preferentially shunted towards complex lipid synthesis and palmitoyl-CoA is used in *de novo* sphingolipid synthesis whereas both are substrates for acyl chain elongation and ceramide synthesis through the action of ELOVL and CerS, respectively. In agreement with this model, we found that palmitate increased saturated LC and VLC (C16:0-C24:0) dihydroceramides and ceramides as well as dihydrosphingomyelins, sphingomyelins and glucosylceramides. Using labelled serine, we observed an increase in ceramide flux characterized by a significant rise in total amount of labelled dihydroceramides and ceramides in response to palmitate. These results are consistent with previous studies describing an increase in ceramide flux in palmitate-exposed MIN6 cells (48). In contrast to palmitate, oleate did not change global sphingolipid flux or steady-state levels. Instead, oleate specifically increased the synthesis of unsaturated C24:1 sphingolipids including dihydroceramides, ceramides, dihydrosphingomyelins, sphingomyelins and glucosylceramides. Although oleate does not affect levels of ceramide species in MIN6 cells (11), to our knowledge, the effect of oleate on the synthesis of other sphingolipid species in the β cell had not been previously described.

Knockdown of ELOVL1 or ACBP in rat islets inhibited oleate-induced β-cell proliferation. By elongating SFA and MUFA to generate C24:0 and C24:1 acyl-CoA, ELOVL1 controls the synthesis of VLC sphingolipids (33). ELOVL1 knockdown in HeLa cells and knockout in mice dramatically reduces C24 sphingolipid levels (19,49). By binding acyl-CoA, modulating expression of genes involved in FA metabolism (50) and interacting with CerS and ELOVL1, ACBP regulates VLC sphingolipid synthesis (32). Accordingly, loss of ACBP in yeast abrogates FA elongation and reduces sphingolipid synthesis (31) and ACBP knockout mice exhibit a significant decrease in VLC acyl-CoA (32). In view of the effect of oleate on sphingolipid metabolism described above, we conclude that C24:1 sphingolipids contribute to oleate-induced β-cell proliferation. Interestingly, unlike oleate, palmitoleate, an n-7 MUFA not elongated into VLC acyl-CoA by ELOVL (33), did not induce β-cell proliferation. We conclude that oleoyl-CoA is likely among a limited number of FA-CoA species that provide the necessary substrate for ELOVL1 to increase the pool of C24:1 acyl-CoA leading to the synthesis of C24:1 sphingolipids and β-cell proliferation.

In conclusion, our results show that oleate-induced β-cell proliferation in isolated rat islets requires the synthesis of C24:1 sphingolipids that is regulated by the availability of oleate. Our findings contribute to the understanding of chain length-specific properties of sphingolipids and support a role for a FA-CoA-C24:1 sphingolipid axis in adaptative β-cell mass expansion in the face of nutrient excess.

## ACKNOWLEDGEMENTS

We thank Caroline Tremblay from the CRCHUM for valuable technical assistance; Grace Fergusson and Mélanie Éthier from the Rodent Metabolic Phenotyping core of the CRCHUM for assistance with *in vivo* studies and islet isolations; Julien La Montagne and Alexia Grangeon from the Metabolomics core of the CRCHUM for S1P and SKI II measurements; and Dominique Gauchat and Philippe St-Onge from the Flow Cytometry core of the CRCHUM for assistance with measurements of β-cell proliferation.

## AUTHOR CONTRIBUTION

A.L.C. Conceptualization, Methodology, Investigation, Formal Analysis, Writing – Original Draft; A.V. Methodology, Investigation, Formal Analysis; T.S.T. Investigation, Formal Analysis; I.R, V.S.M. and M.R. Investigation; J.G. Conceptualization, Validation, Writing – Review and Editing, Supervision; C.D.R, W.L.H, S.A.S, V.P Conceptualization, Validation, Writing – Review and Editing, Supervision, Funding Acquisition, Project Administration. V.P. is the guarantor of this work and, as such, had full access to all the data in the study and takes responsibility for the integrity of the data and the accuracy of the data analysis.

## PRIOR PRESENTATION

Parts of this study were presented at the 79th Scientific Sessions of the American Diabetes Association, San Francisco, CA, 7–11 June 2019.

