## Supplementary Materials for "Very Long-Chain Unsaturated Sphingolipids Mediate Oleate-Induced Rat β-Cell Proliferation"

**SUPPLEMENTARY RESEARCH DESIGN AND METHODS**

**Reagents and Solutions**

RPMI-1640, MEM, FBS and dialysed FBS were from Life Technologies Inc. (Burlington, ON, Canada). Penicillin/Streptomycin was from Multicell Wisent Inc. (Saint-Jean-Baptiste, QC, Canada). FA-free BSA was from Equitech-Bio (Kerrville, TX, USA). Oleate and palmitate were from Sigma Aldrich (Saint-Louis, MI, USA), linoleate from MP Biomedicals (Solon, OH, USA) and palmitoleate, Myriocin and SKI II from Cayman Chemical (Ann Arbor, MI, USA). Adenoviruses expressing shRNA against SphK1 (Adv-shSphK1), SphK2 (Adv-shSphK2), ELOVL1 (Adv-shELOVL1), ACBP (Adv-shACBP) and control scrambled shRNA (Adv-shCTL) were from Vector Biolabs (Malvern, PA, USA).

**Immunostaining of Pancreatic Sections**

Pancreata were fixed for 4 h in 4% paraformaldehyde (PFA) and cryoprotected overnight in 30% sucrose. Tissue was then embedded in optimal cutting temperature (OCT) compound, frozen, sectioned at 8 μm, and mounted on Superfrost Plus slides (Life Technologies Inc., Burlington, ON, Canada). Antigen retrieval was performed using sodium citrate buffer (pH=6). B-cell proliferation was measured by immunohistochemical staining for Ki67 and insulin (Ins) or Nkx6.1 as described (1). Primary antibodies and dilutions are listed in Supplementary Table 1. Secondary antibodies were from Jackson ImmunoResearch (West Grove, PA, USA). Images were acquired with a fluorescence microscope (Zeiss, Thornwood, NY). B-cell proliferation was expressed as a percentage of double-positive Ki67^+^ and Ins^+^ (or Nkx6.1^+^) cells over the total Ins^+^ (or Nkx6.1^+^) cells. At least 1,500 β-cells were manually counted per condition. The experimenter was blind to group assignments.

**Plasma SKI II Levels in Nutrient-infused Rats**

SKI II was extracted from rat plasma by protein precipitation. Briefly, 150 µl of acetonitrile was added to 50 µl of rat plasma, vortexed and incubated on ice for 10 min. Samples were then centrifuged 16,000 x g for 10 min at 4°C and the supernatants evaporated under a stream of nitrogen. Samples were re-suspended in 50 µl acetonitrile prior to analysis by LC-ESI-MS/MS according to a method modified from French et al. (2). MRM analysis was performed in positive ion mode to obtain spectra for SKI II 303/174.

**Beta-cell Proliferation and Apoptosis in Isolated Islets**

To assess β-cell proliferation in isolated islets by immunohistochemistry, following treatment islets were embedded in OCT compound, frozen, sectioned at 8 μm, and mounted on Superfrost Plus slides. Antigen retrieval was performed using sodium citrate buffer (pH=6) after 30 min fixation in 3% PFA. B-cell proliferation was measured by immunohistochemical staining for Ki67 and Ins as mentioned above for pancreatic sections.

To assess β-cell proliferation by flow cytometry, at the end of the treatment islets were dispersed in Accutase (1 μl/islet; Innovative Cell Technologies Inc., San Diego, CA) for 10 min at 37° C dead cells labeled using the LIVE/DEAD™ Aqua (405 nm) or LIVE/DEAD^TM^ Red (488 nm) Dead Cell Stain Kit (BD Bioscience). EdU detection, using Click-iT Plus EdU Alexa Fluor 488 or Click-iT Plus Pacific Blue Flow Cytometry Assay Kit, and immunostaining was performed according to the manufacturer’s instructions (Thermo Fisher Scientific, Waltham, MA). Fluorophore-coupled primary antibodies and dilutions are listed in Supplementary Table 1. Flow cytometric analysis was carried out using a LSRIIB flow cytometer with FACSDiva software (BD Biosciences). Data was analyzed using FACSDiva or FlowJo v10.7 software (Ashland, OR; <https://www.flowjo.com/solutions/flowjo>). Dead-cell stain, EdU, C-peptide (Cpep), Ins and glucagon (GCG) labelled cells were detected using the 405-, 488-, 640- and 561-nm lasers coupled with 525/50-, 530/30-, 670/14- and 586/15-nm BP filters, respectively. Proliferation was calculated as the percentage of double-positive EdU^+^ and Cpep^+^ (or Ins^+^) cells over the total Cpep^+^ (or Ins^+^) cell population. At least 10,000 Cpep^+^ (or Ins^+^) cells were counted in each sample.

To measure β-cell proliferation after adenoviral infection we used the GFP reporter to identify GFP positive (GFP^+^) and negative (GFP^-^) cells within the β-cell population. Proliferation was expressed as the percentage of Cpep^+^ (or Ins^+^), GFP^+^ and EdU^+^ cells over the double-positive Cpep^+^ (or Ins^+^) and GFP^+^ cells. At least 2,000 Cpep^+^ (or Ins^+^) GFP^+^ cells were counted in each sample when islets were infected with 1 PFU and at least 5,000 Cpep^+^ (or Ins^+^) GFP^+^ cells were counted when islets were infected with 10 or 100 PFU.

To assess β-cell apoptosis by flow cytometry, at the end of the treatment islets were dispersed as described above, washed twice with PBS. Dead cells were labeled using eBioscience Fixable Viability Dye eFluor 506 and 780 (Thermo Fisher Scientific) and apoptotic cells detected using the eBioscience Annexin V Apoptosis Detection Kit FITC (Thermo Fisher Scientific). Samples where then fixed, permeabilized and stained for Cpep according to manufacturer’s instruction (Thermo Fisher Scientific). Fluorophore-coupled primary antibodies and dilutions are listed in Supplementary Table 1, respectively. Dead-cell stain, Cpep, Annexin V labelled cells were detected using the 405- and 640-, 640-, 488- lasers coupled with 525/50-, 780/60-, 670/14-, 530/30-nm BP filters, respectively. Apoptosis was calculated as the percentage of Cpep^+^/Annexin V^+^/eFluor506^-^ cells over the total Cpep^+^ cells. At least 20,000 Cpep^+^ cells were counted in each sample.

**Quantitative RT-PCR**

To assess SphK knockdown efficiency, SphK1 and SphK2 expression levels were measured by quantitative RT-PCR of infected GFP^+^ cells. Briefly, at the end of treatment, islets were dissociated in Accutase (1 μl/islet; Innovative Cell Technologies Inc., San Diego, CA) for 10 min at 37°C, washed with cold PBS, resuspended in PBS+1% BSA and passed through a 50-μm filter prior to flow cytometric sorting using a FACSAria II flow cytometer with FACSDiva software (BD Biosciences, San Jose, CA). To assess ELOVL1 and ACBP knockdown efficiency gene expression was measured by quantitative RT-PCR in whole islets. Total RNA was extracted from 150 to 200 whole islets or at least 50,000 sorted β-cells using the RNeasy Micro kit (Qiagen, Valencia, CA). RNA was quantified by spectrophotometry using a NanoDrop 2000 (Life Technologies Inc.), and 1 μg of RNA was reverse transcribed. Real-time PCR was performed using the Rotor-Gene SYBR Green PCR kit (Qiagen). Results are expressed as the ratio of target mRNA to cyclophilin A RNA levels and normalized to the levels in control islets. Primers sequences are listed in Supplementary Table 2.

**Sphingolipidomic Analyses**

For S1P measurement (Fig. 1E), batches of 200 islets were washed in cold PBS and suspended in 2 ml methanol, containing 5 nM C17-S1P (Avanti Polar Lipids) as internal standard, followed by 1 ml of chloroform and 0.5 ml of 0.1 M HCl in water. Samples were vortexed, incubated for 10 min at room temperature and centrifuged at 4500 x g for 10 min. The lipid-containing lower phase was collected and dried under a stream of nitrogen. Samples were resuspended in 50 µl MeOH prior to analysis by LC-ESI-MS/MS according to a method modified from Berdyshev et al. (3). MRM analysis was performed in negative ion mode to obtain spectra for AcS1P-C17 490/448 and AcS1P-C18 504/462.1.

Sphingolipidomic analysis (Fig. 4B, C & 7) was conducted using methods described previously (4). Lipid were extracted from isolated rat islets (200 islets each) and separated on an Acquity UPLC CSH C18 1.7 µm 2.1 x 50 mm column maintained at 60°C connected to an Agilent HiP 1290 Sampler, Agilent 1290 Infinity pump, and Agilent 6490 triple quadrupole (QqQ) mass spectrometer. Sphingolipids are detected using dynamic multiple reaction monitoring (dMRM) in positive ion mode. Source gas temperature is set to 210°C, with a gas (N_2_) flow of 11 L/min and a nebulizer pressure of 30 psi. Sheath gas temperature is 400°C, sheath gas N_2_ flow of 12 L/min, capillary voltage is 4000 V, nozzle voltage 500 V, high pressure RF 190 V and low-pressure RF is 120 V. Injection volume is 2 µL and the samples are analyzed in a randomized order with the pooled QC sample injected eight times throughout the sample queue. Mobile phase A consists of ACN: H_2_O (60:40 v/v) in 10 mM ammonium formate and 0.1% formic acid, and mobile phase B consists of IPA:ACN:H_2_O (90:9:1 *v/v*) in 10 mM ammonium formate and 0.1% formic acid. The chromatography gradient starts at 15% mobile phase B, increases to 30% B over 1 min, increases to 60% B from 1-2 min, increases to 80% B from 2-10 min, and increases to 99% B from 10-10.2 min where it’s held until 14 min. Post-time is 5 min and the flowrate is 0.35 mL/min throughout. Collision energies and cell accelerator voltages were optimized using sphingolipid standards with dMRM transitions as [M+H]^+^→[*m/z* = 284.3] for dihydroceramides, [M+H]^+^→[*m/z* = 287.3] for isotope labeled dihydroceramides, [M-H_2_O+H]^+^→[*m/z* = 264.2] for ceramides, [M-H_2_O+H]^+^→[*m/z* = 267.2] for isotope labeled ceramides and [M+H]^+^→[M-H_2_O+H]+ for all targets. Results from liquid chromatography-mass spectrometry (LC-MS) experiments are collected using Agilent Mass Hunter Workstation and analyzed using the software package Agilent Mass Hunter Quant B.07.00. Sphingolipids are quantitated based on peak area ratios to the standards added to the extracts.

Lipid extraction, LC-MS analysis and data processing (Fig. 5) were done as previously described (5). In brief, lipids were extracted from isolated rat islets (500 islets each) and spiked with six internal standards: LPC 13:0, PC19:0/19:0, PC14:0/14:0, PS12:0/12:0, PG15:0/15:0 and PE17:0/17:0 (Avanti Polar Lipids Inc, Alabaster, USA). Samples were injected (2μL and 4 μL in positive and negative ionization mode respectively) into a 1290 Infinity High-pressure liquid chromatography (HPLC) coupled with a 6530 Accurate Mass Quadrupole Time-of-Flight (Q-TOF) (Agilent Technologies Inc., Santa Clara, USA) via a dual electrospray ionization (ESI) source. Elution of lipids was assessed on a Zorbax Eclipse plus column (C18, 2.1 x 100 mm, 1.8 μm, Agilent Technologies Inc.) maintained at 40°C using an 83 min chromatographic gradient of solvent A (0.2% formic acid and 10 mM ammonium formate in water) and B (0.2% formic acid and 5 mM ammonium formate in methanol/acetonitrile/methyl tert-butyl ether [MTBE], 55:35:10 [v/v/v]). A list of MS features, characterized by mass, retention time and signal intensity (area under the curve), was extracted using Mass Hunter B.06.00 (Agilent Technologies Inc.). Subsequent data mining was achieved using an in-house script that applies alignment of the chromatographic runs, frequency filtering (80 % in one condition), signal intensity normalization using cyclic loess algorithm and imputation of missing values with k-nearest neighbor on scaled data. MS signals for 28 sphingolipids including their acyl side chains (20 sphingomyelins (SM), 6 ceramides (Cer), 4 hexoceramides (GlcCer)) were extracted using an in-house database. Data were expressed as normalized signal intensity.

**SUPPLEMENTARY TABLES**

**Supplementary Table 1**. Sources of antibodies used

| Antibody | Dilution/Concentration | Company | Cat # |
| --- | --- | --- | --- |
| **Immunohistochemistry** |  |  |  |
| Insulin | 1:500 | DAKO | A0564 |
| Nkx6.1 | 5μg/ml | DSHB | F55A12 |
| Ki67 | 1:500 | Abcam | 15580 |
| **Flow cytometry** |  |  |  |
| Alexa Fluor® Mouse anti-C-peptide | 1:25 | BD Biosciences | 565831 |
| Alexa Fluor® Mouse anti-insulin | 1:25 | BD Biosciences | 565689 |
| PE Mouse Anti-Glucagon | 1:25 | BD Biosciences | 565860 |

**Supplementary Table 2**. RT-PCR primers.

| Primer name | Forward | Reverse |
| --- | --- | --- |
| Cyclophilin | CTTGCTGCAGACATGGTCAAC | GCCATTATGGCGTGTGAAGTC |
| Sphk1 | AGATGGATGAGAGGGAGGGT | CCAAGTGCACCCAAACTACC |
| Sphk2 | GGCAGTTAACCATCATGGCG | GAGGCTAGCGTCACAGAGAG |
| ELOVL1 | GGCCCTGATCCCTTTGAACC | CGGGGATCTGCACACTTCAT |
| ACBP | AGCTGAAAGGAACTTCCAAG | GGTATTATGTCACACATGTGGC |

**SUPPLEMENTARY FIGURES**

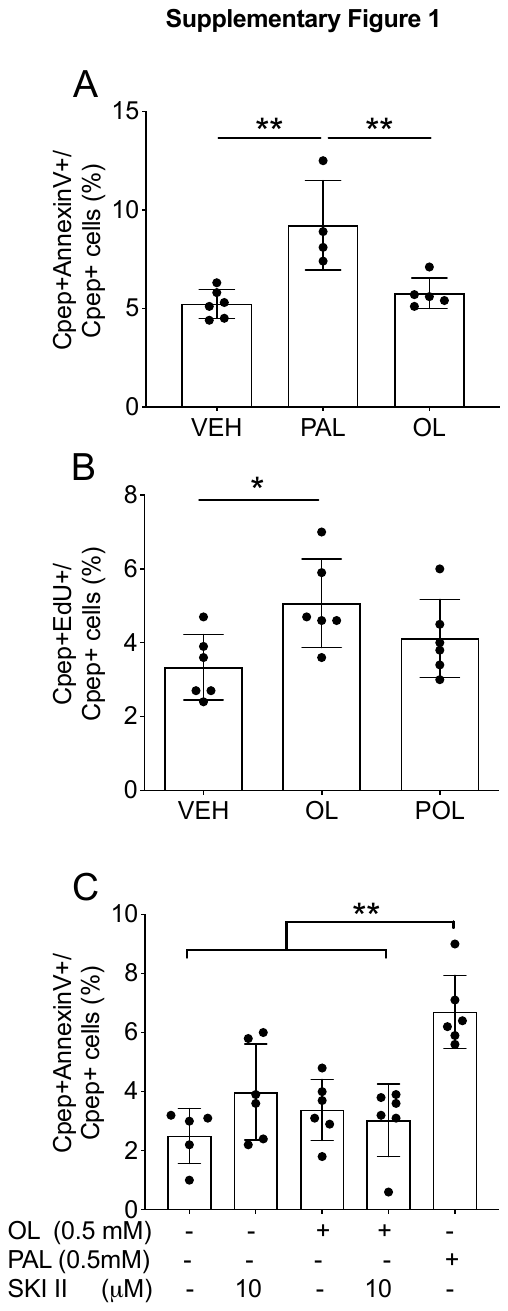

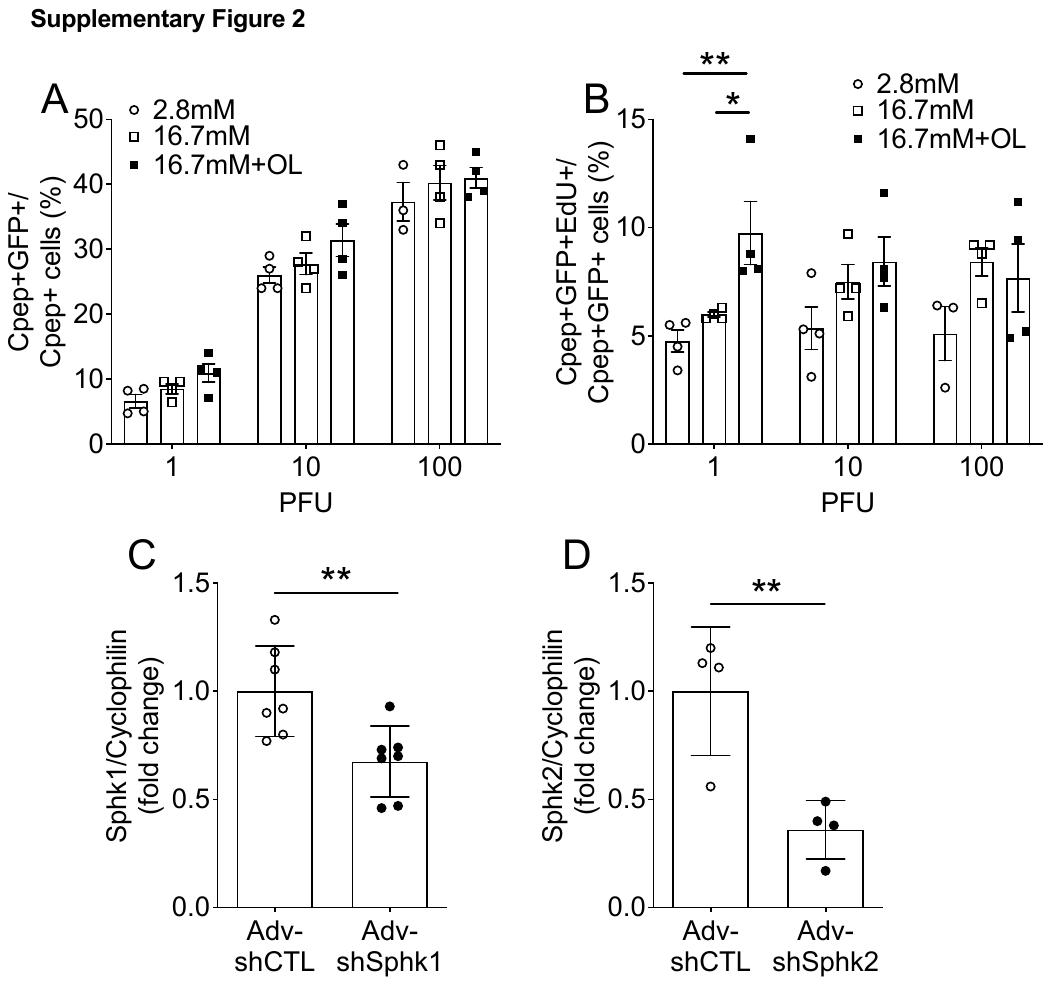

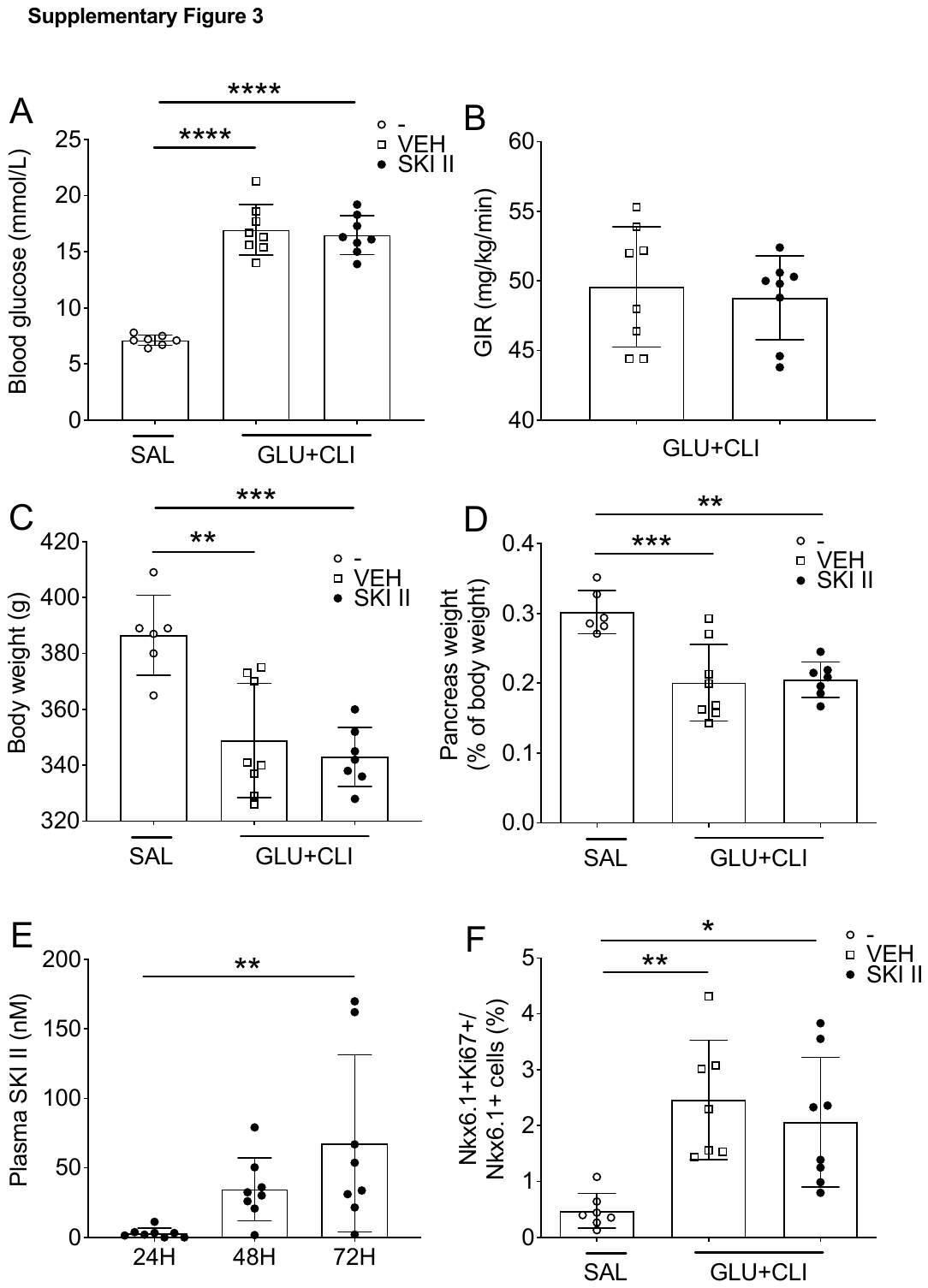

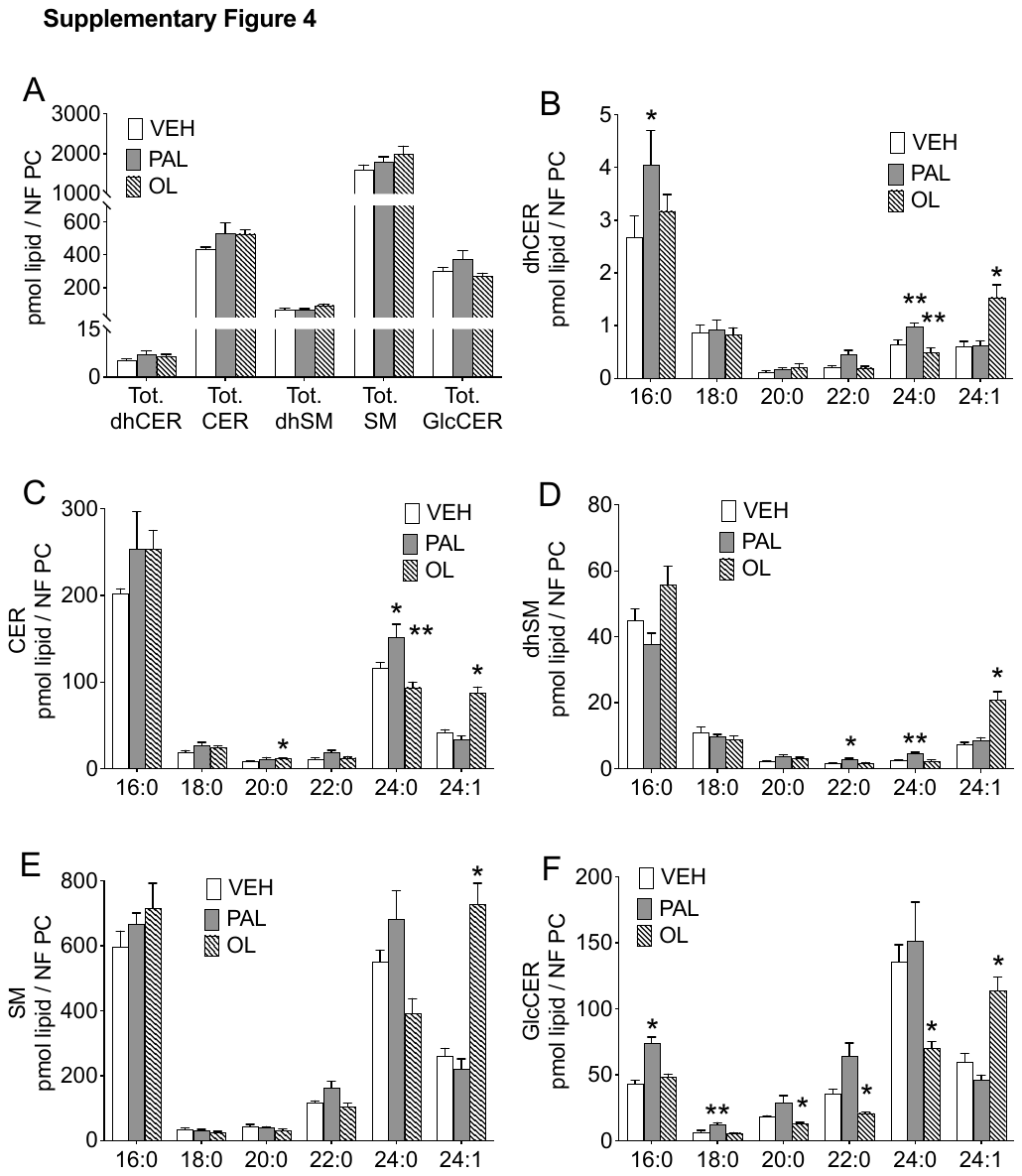

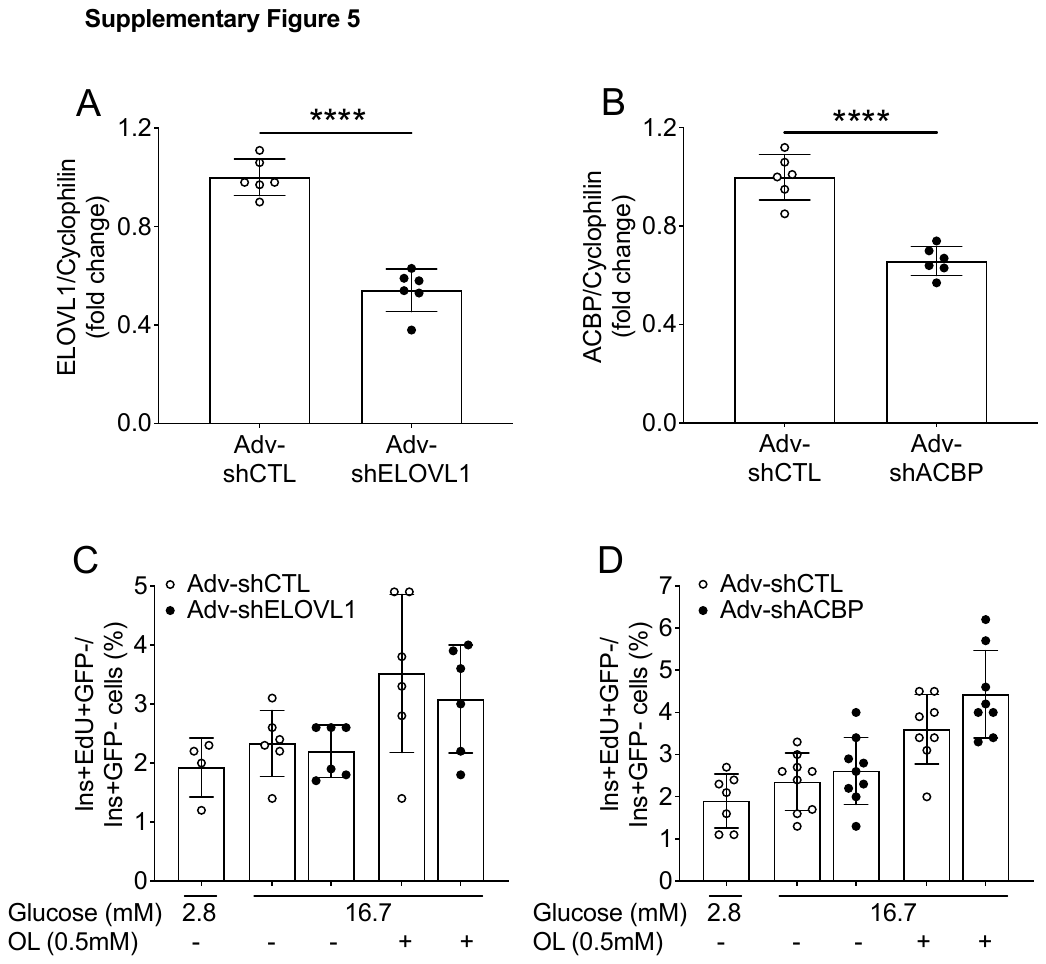

**SUPPLEMENTARY FIGURE LEGENDS**

**Supplementary Figure 1. B-cell** **proliferation and apoptosis in response to fatty acids in isolated rat islets**. (A-C) Isolated rat islets were exposed to 16.7 mM glucose and palmitate (PAL, 0.5 mM), palmitoleate (POL, 0.5 mM), oleate (OL, 0.5 mM) or vehicle (VEH) with or without SKI II (10 μM) for 48 h as indicated. (A, C) β-cell apoptosis was assessed by flow cytometry following staining for AnnexinV and Cpep and presented as a percentage of AnnexinV^+^/Cpep^+^ over total Cpep^+^ cells. (B) β-cell proliferation was assessed by flow cytometry following staining for EdU and C-peptide (Cpep) and presented as a percentage of EdU^+^/Cpep^+^ over total Cpep^+^ cells. (B, D) Data represent individual values and are expressed as means +/- SEM. *p<0.05, **p<0.01 following one-way ANOVA with Tukey’s multiple comparisons test.

**Supplementary Figure** **2.** **Adenoviral-mediated knockdown of SphK.** (A, B) Isolated rat islets were infected with 1, 10 and 100 PFU of Adv-shCTL in the presence of 2.8 or 16.7 mM glucose with or without oleate (OL) (*n*=3-4). (A) The level of adenoviral transduction of β-cells was assessed by flow cytometry following staining for C-peptide (Cpep) and expressed as a percentage of GFP^+^/Cpep^+^ over total Cpep^+^ cells. (B) β-cell proliferation was assessed by flow cytometry following staining for EdU and Cpep and expressed as percentage of EdU+/GFP^+^/Cpep^+^ over total GFP^+^/Cpep^+^ cells. (C, D) Sphk1 and Sphk2 mRNA levels were determined on FACS-sorted GFP^+^ cells following 48 h-infection with Adv-shCTL or Adv-shSphk1 (C) or Adv-shSphk2 (D) (*n*=4-7). mRNA was quantified by RT-PCR and normalized to cyclophilin. Data are presented as the fold change over the control condition (Adv-shCTL). Data represent individual values and are expressed as means +/- SEM. *p<0.05, **p<0.01 following one-way ANOVA with Tukey’s multiple comparisons test (A, B) or Student’s t-test as compared with Adv-shCTL (C, D).

**Supplementary Figure** **3.** **Metabolic parameters and β-cell proliferation in SKI II treated nutrient-infused rats.** 2-month-old Wistar rats were infused with saline (SAL, white round) or glucose and ClinOleic (GLU+CLI) for 72 h. GLU+CLI-infused animals received either vehicle (VEH, white square) or SKI II by oral gavage (50mg/kg/d, black round) (*n*=7-8). (A-D) Average blood glucose (A) and GIR (B), measured during the infusion, and body (C) and pancreas weight (D) at the end of the infusion. (E) Plasma SKI II levels at 24, 48 and 72 h in GLU+CLI-infused animals. (F) β-cell proliferation was assessed by immunostaining for Ki67 and Nkx6.1 and presented as a percentage of Ki67^+^/Nkx6.1^+^ cells over total Nkx6.1^+^ cells. Data represent individual values and are expressed as means +/- SEM. *p<0.05, **p<0.01, ***p<0.005, ****p<0.001 following one-way ANOVA with Tukey’s multiple comparisons test (A, C-F) or following Student’s t-test as compared with the VEH group (B).

**Supplementary Figure 4: Sphingolipid profile of isolated rat islets exposed to palmitate or oleate.** Isolated rat islets were exposed to 16.7 mM glucose and palmitate (PAL, 0.5 mM), oleate (OL, 0.5 mM) or vehicle (VEH) for 48 h. Lipidomic analyses were performed by LC-MS/MS on islets extracts at the end of the treatment. Levels of total sphingolipids (A) and dihydroceramides (dhCER) (B), ceramides (CER) (C), dihydrosphingomyelins (dhSM) (D), sphingomyelins (SM) (E) and glucosylceramides (GlcCER) (F) grouped according to acyl-chain length are expressed in pmol normalized to phosphatidyl choline species. Data represent means +/- SEM (*n*=4). *p<0.05, **p<0.01, following one-way (A) or 2-way ANOVA (B-F) with Dunett’s multiple comparisons test as compared to the VEH condition.

**Supplementary Figure** **5:** **Adenoviral-mediated knockdown of ELOVL1 and ACBP in isolated rat islets.** (A, B) ELOVL1 and ACBP mRNA were measured following 48h-infection with Adv-shCTL, Adv-shELOVL1 (A) or Adv-shACBP (B) in isolated rats islets (*n*=6). mRNA was quantified by RT-PCR and normalized to cyclophilin. Data are presented as the fold change over the control condition (Adv-shCTL). (C, D) Isolated rat islets were infected with Adv-shCTL, Adv-shELOVL1 (C) or Adv-shACBP (D) and exposed to 2.8 or 16.7 mM glucose with or without oleate (OL, 0.5 mM) as indicated. Proliferation of uninfected (GFP^-^) β cells was assessed by flow cytometry following staining for EdU and insulin (Ins) and presented as a percentage of EdU^+^/Ins^+^/GFP^-^ over total Ins^+^/GFP^-^ cells. Data represent individual values and are expressed as means +/- SEM. ****p<0.001 following Student’s t-test (A-B) or Mixed effect analysis with Sidak’s multiple comparisons test as compared with Adv-shCTL condition (C, D).
